# Epigenetic regulation of innate immune genes and enhanced interleukin-10 expression underlie chronic subclinical *Plasmodium chabaudi* infection

**DOI:** 10.1101/2023.02.23.529826

**Authors:** Leandro de Souza Silva, Yen Anh H. Nguyen, Brian G. Monks, Catherine S. Forconi, Juliet N. Crabtree, Tomás Rodriguez, Nelsy De Paula Tamburro, Erik J. Sontheimer, Gabor L. Horvath, Zeinab Abdullah, Eicke Latz, Daniel R. Caffrey, Evelyn A. Kurt-Jones, Ricardo T. Gazzinelli, Katherine A. Fitzgerald, Douglas T. Golenbock

## Abstract

Subclinical (asymptomatic) parasitemia is very common amongst *Plasmodium*-infected individuals. The immunological mechanisms underlying subclinical parasitemia remain elusive. We investigated the immune regulatory mechanisms behind chronic asymptomatic *Plasmodium* infection using mice lacking humoral immunity (µMT^−/−^ mice). µMT^−/−^ mice became chronically infected, despite lacking outward signs of disease, and exhibited increased macrophage numbers, decreased dendritic and CD4 cells, massive hemozoin accumulation in the spleen and bone marrow, and inadequate hematopoiesis. These changes were accompanied by high circulating levels of interleukin-10 (IL-10), enhanced chromatin accessibility of the STAT3 promoter, and enhanced STAT3 binding to the IL-10 promoter in macrophages. Inhibition of IL-10 signaling, despite promoting parasite clearance, resulted in a proinflammatory response, weight loss, and mortality. These results suggest that epigenetic changes induced by chronic *P. chabaudi* infection lead to high levels of circulating IL-10, protecting chronically infected mice against an excessive inflammatory response to high levels of blood-stage parasites.

**Author summary:** Malaria is a life-threatening disease with a range of symptoms, and it is induced in humans by infections with different species of *Plasmodium*. Highly prevalent in endemic regions, asymptomatic *Plasmodium* infections are related to long-term exposure to the parasite due to multiple infections and have been demonstrated in human and mouse studies to be associated with elevated levels of IL-10. However, how IL-10 levels remain elevated in the circulation in individuals over the long term has not been determined. We used a mouse model of chronic asymptomatic *Plasmodium* infection to investigate the mechanisms by which IL-10 levels are elevated during chronic asymptomatic infection. Our results show that epigenetic changes in immune genes of myeloid origin could be responsible for the elevated levels of IL-10, and that IL-10 signaling protected chronically infected mice from a severe inflammatory response induced by the infection.

## Introduction

In 2020, *Plasmodium* spp. infections were responsible for ∼241 million cases of malaria and 627,000 deaths worldwide (WHO, 2021). Upon infection, the spectrum of clinical outcomes ranges from life-threatening syndromes (*e.g.*, severe anemia and cerebral malaria) to an uncomplicated febrile disease to asymptomatic infection. The exact clinical syndrome that accompanies an infection appears to depend on a complex interaction between factors associated with the parasite, the environment, and the human host (Miller et al., 2002).

Asymptomatic malaria is more accurately referred to as “subclinical malaria” because subtle symptoms and chronic health effects may occur but not lead to clinical diagnosis and treatment. Asymptomatic infections may present with a wide range of parasitemia levels, from submicroscopic to microscopic levels that appear to exceed the putative pyrogenic threshold (Chen et al., 2016). Asymptomatic *Plasmodium* spp. infections are especially prevalent in areas of high endemicity and are related to chronic exposure to the parasite through repeated infections (Roucher et al., 2012). This naturally acquired “immunity” protects the host against severe disease but only partially controls the parasitic burden (Riley et al., 2006) and does not protect against new infections (Gatton and Cheng, 2004). Studies in both humans and mice have shown a role for T cell-derived interleukin-10 (IL-10) in this process (Couper et al, 2008; Freitas do Rosário et al 2012; Kumar et al., 2019). However, how elevated circulating IL-10 levels are maintained upon chronic exposure to the *Plasmodium* spp. parasite is still not well understood.

IL-10 is a key regulatory cytokine that protects against severe *Plasmodium*-induced pathology during acute infections. A study of subclinical malaria in B-cell-deficient mice (J_H_^−/−^) infected for up to 300 days with *P*. *chabaudi adami* showed high circulating IL-10 levels (Weidanz et al., 2005). In humans, the role of IL-10 is increasingly under investigation as a major cytokine responsible for subclinical malaria. Individuals living in malaria hyperendemic areas are repeatedly exposed to the parasite over the course of many years. The proinflammatory response to subsequent infections typically decreases with age (Tran et al., 2013). A study of children from an area hyperendemic for *P. falciparum* demonstrated that the frequency of IL-10+/IFNγ+ CD4+ T cells positively correlates with the cumulative number of malaria episodes (Jagannathan et al., 2014). In addition, purified monocytes from adults in endemic areas have increased IL-10 production after *in vitro* exposure to *P. falciparum-*infected erythrocytes compared to monocytes from individuals who were never exposed to the parasite (Guha et al., 2021). IL-10 production upon challenge with *P. falciparum-*infected erythrocytes also increased with age starting in childhood (Guha et al., 2021). Although studies clearly support a role for IL-10 in chronic asymptomatic infections, the mechanisms of IL-10 during this process and how high circulating levels of IL-10 are maintained over time have not been examined. We hypothesize that epigenetic changes induced by the constant presence of the parasite drive persistent circulating levels of IL-10 that control the inflammatory response against the blood stage of the parasite.

In this study, we used B6.129S2-*Ighm^tm1Cgn^*/J mice (hereafter referred to as μMT^−/−^ mice), a strain of mice that lack mature B cells. The μMT^−/−^ mice were infected with *P*. *chabaudi* AS to model chronic subclinical malaria. We observed that μMT^−/−^ mice became chronically infected but had no external signs of disease. In contrast, wild-type C57BL/6J mice always clear *P. chabaudi* infection. Despite the lack of apparent disease, chronically infected μMT^−/−^ mice had suppressed hematopoiesis, altered immune cellular distribution, and massive hemozoin accumulation in splenic tissue, accompanied by high circulating levels of IL-10 compared to WT mice. Furthermore, chronic infection increased the chromatin accessibility of STAT3 gene promoter in bone marrow-derived macrophages from μMT^−/−^ compared to WT mice. Chronically infected-μMT^−/−^ mice had enhanced binding of STAT3 to the IL-10 promoter. Using neutralizing antibodies to block IL-10 signaling in chronically infected mice resulted in a resurgent inflammatory response accompanied by increased circulating neutrophils and monocytes, as well as weight loss, a decline in physical activity, and death. Our data suggest that high circulating IL-10 levels, maintained by epigenetic changes, promote asymptomatic parasitemia by suppressing the inflammatory response through the IL-10R/STAT3 signaling axis.

## Results

### *Plasmodium chabaudi*-infected μMT^−/−^ mice have patent chronic infection and defective hematopoiesis

To investigate the mechanisms of chronic subclinical malaria, we established *Plasmodium chabaudi*-infected μMT^−/−^ mice as a model for subclinical (often referred to as asymptomatic) malaria (S1 Fig). C57BL/6J (WT) and μMT^−/−^ mice were infected with the *P. chabaudi* AS strain expressing green fluorescent protein (GFP). Infection was monitored by flow cytometry and PCR of peripheral blood (S2 Fig). Further analyses were carried out at various phases of the infection referred to as denoted in Fig. S1. Upon *P. chabaudi* infection, both WT and μMT^−/−^ mice had a peak of parasitemia ranging from ∼15– 35% of red blood cells (RBCs) between 7 and 10 days post-infection (p.i.), consistent with the results of others (Aiello and Caffrey, 2012; Stephens et al., 2012). In WT mice, the parasitemia typically dropped to sub-patent levels (chronic phase) by day 20 p.i. In contrast, infected μMT^−/−^ mice showed persistent patent parasitemia for more than 3 months p.i.; (Fig 1A). Indeed, in our several years of using the μMT^−/−^ model, we have never observed a single μMT^−/−^ mouse spontaneously clear an infection. Thus, *P. chabaudi* infection of μMT^−/−^ mice enables studies of persistent parasitemia. After 99 days p.i., when GFP+ erythrocytes were consistently undetectable in blood samples of infected WT mice (Fig 1A), nested PCR confirmed the absence of parasitemia (Fig 1B). Persistently infected μMT^−/−^ mice had a low percentage of mature RBCs (Fig 1C) and very high levels of reticulocytes (Fig 1D) in the peripheral blood. In addition, bone marrow from chronically infected μMT^−/−^ mice had lower levels of mature RBCs (Fig 1E) and higher levels of reticulocytes (Fig 1F) compared to controls, consistent with a premature release of immature RBCs due to ongoing hemolysis.

**Fig 1.**
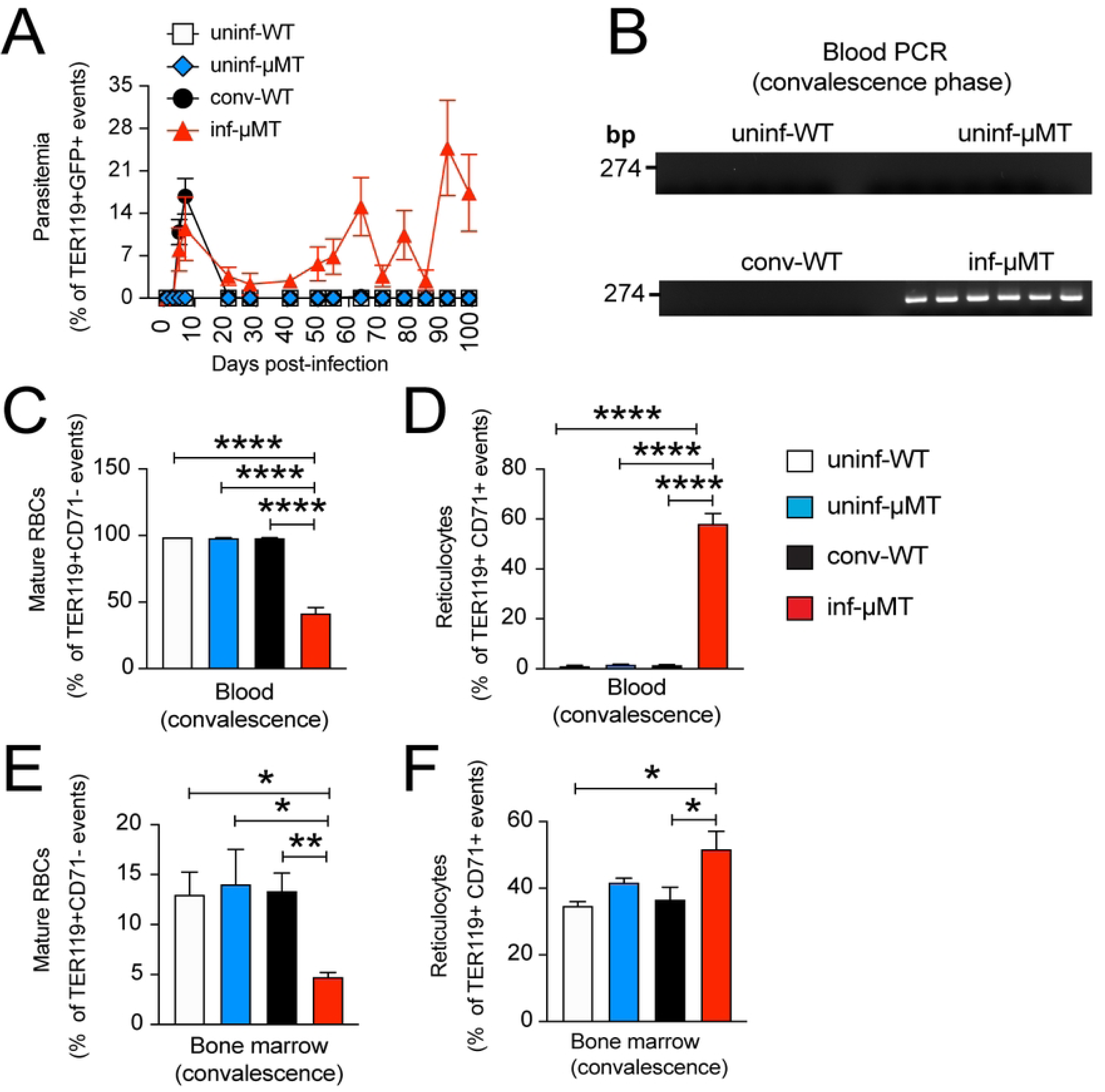
*Plasmodium chabaudi*-infected μMT^−/−^ mice have persistent patent parasitemia accompanied by defective hematopoiesis. (A) Parasitemia of convalescent WT (black circle, n=6) and μMT^−/−^ (red triangle, n=6) mice infected with fluorescent *P*. *chabaudi* AS are shown as a percentage of mature red blood cells (RBCs) defined as TER119+ cells (10^6^ RBCs were counted per sample). Blood samples of uninfected WT (white square, n=6) and μMT^−/−^ (blue diamond, n=6) mice were used as negative controls. (B) PCR amplification of blood samples from uninfected WT, uninfected μMT^−/−^, infected WT (convalescent), and chronically infected μMT^−/−^ mice 99 days post-infection (dpi). (C) Percentage of mature RBCs and (D) reticulocytes (defined as TER119^+^CD71^+^ events) from mouse peripheral blood 99 dpi, n=3. (E) Percentage of mature RBCs and (F) reticulocytes from the bone marrow of uninfected WT, uninfected μMT^−/−^, infected WT (convalescent), and chronically infected μMT^−/−^ mice (n=4). Data are shown as mean ± SEM. (C and D) * P<0.05, ** P<0.01, **** P<0.0001, as determined by one-way ANOVA followed by Tukey’s multiple comparisons test or unpaired Student’s t-test (E and F).

### Chronically infected μMT^−/−^ mice appear to be asymptomatic

Low levels of physical activity are commonly observed during the acute phase (days 7– 10 p.i.) of *P. chabaudi* infection in C57BL/6J mice. These mice exhibit weight loss, hypothermia, and other signs of systemic inflammation (Franklin et al., 2007; Wilson et al., 2016). Observable signs of infection in WT mice disappear after the acute phase, when the parasitemia is reduced to sub-patent levels (chronic phase) until finally, the host completely clears the parasite from the circulation (convalescence phase), a process that could take 30-90 days (Pérez-Mazliah et al. 2019). To investigate if chronically infected μMT^−/−^ mice are truly asymptomatic and thus resemble humans with subclinical malaria, we assessed the clinical status of the infected WT and μMT^−/−^ mice with the use of metabolic cages (TSE Systems). We measured four different parameters covering the three phases of infection in WT mice (acute, chronic, and convalescent). Mice were evaluated for food consumption, water intake, energy expenditure rate, and physical activity. Despite persistent parasitemia, infected μMT^−/−^ mice did not show any discernible signs of disease at any time point (Fig 2) except for the acute phase, at a time point when both infected WT and μMT^−/−^ mice had similar levels of parasitemia (S3 Fig). After the acute phase, no external signs of disease were observed in the infected mice during the chronic and convalescence phases (Figs 2A–2D) despite persistent infection in μMT^−/−^ mice over a seven-week period, a point at which WT mice had cleared their parasitemia (convalescent phase) as demonstrated by PCR (S3 Fig). Hence, long-term infection in μMT^−/−^ mice resembles subclinical malaria in humans.

**Fig 2.**
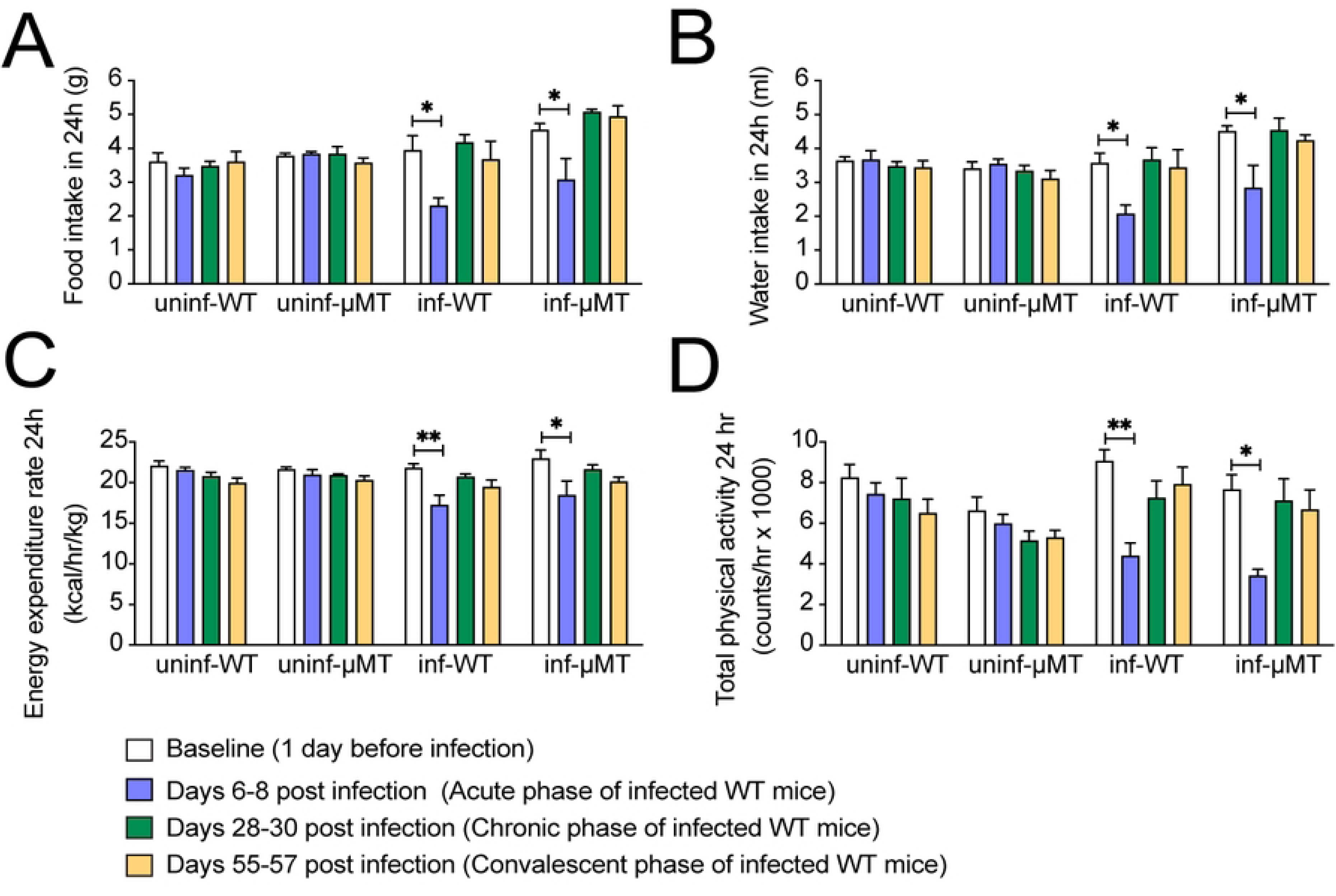
Chronically infected μMT^−/−^ mice have no signs of infection by metabolic cage analysis. (A) Food intake in grams, (B) water intake in mL, (C) energy expenditure rate in kcal/hr/kg, and (D) total physical activity in counts/hr, from uninf-WT, uninf-μMT^−/−^, conv-WT, and inf-μMT^−/−^ mice. Time points were based on the *P. chabaudi* infection profile in WT mice during the acute phase (6–8 dpi), chronic phase (28–30 dpi), and after WT mice have cleared parasitemia as assessed by a negative PCR result (55–57 dpi), referred to as convalescent phase (n=3). Data collected in metabolic cages are expressed as the average of 3 consecutive periods of 24 h. Data are shown as mean ± SEM, *P<0.05, **P<0.01 (compared with baseline values), as determined by unpaired Student’s t-test.

### Chronic infection induces changes in the microarchitecture of the spleen and leads to massive hemozoin accumulation

During the acute phase of infection, *P. chabaudi* induces changes in the splenic microarchitecture by transitory alterations in cellular distribution (Achtman et al., 2003). After the infection resolves, the splenic microarchitecture in WT mice returns to its original form (Achtman et al., 2003). Similar changes were also observed in *P. falciparum* malaria (Urban et al., 2005). To evaluate the kinetics, magnitude, and duration of changes in the splenic microarchitecture in WT and μMT^−/−^ mice, we examined tissue sections by brightfield, reflection confocal microscopy, and immunofluorescence confocal microscopy after recovery from acute symptoms at 99 days p.i. (when infected WT mice were PCR-negative for *P. chabaudi* and thus convalescent while the μMT^−/−^ mice exhibited chronic parasitemia). Uninfected μMT^−/−^ mice had smaller spleens compared with uninfected WT mice, consistent with the lack of mature B cells in these mice. However, chronically infected μMT^−/−^ mice developed massive splenomegaly compared to convalescent WT mice (Fig 3A). Microscopic analysis of tissue sections stained with hematoxylin and eosin (H&E) showed that spleens from uninfected μMT^−/−^ mice have a much smaller white pulp than the WT uninfected control spleens. Convalescent WT mice showed a preserved splenic microarchitecture at 99 days p.i.; however, chronically infected μMT^−/−^ mice showed a complete loss of normal splenic organization (Fig 3B).

**Fig 3.**
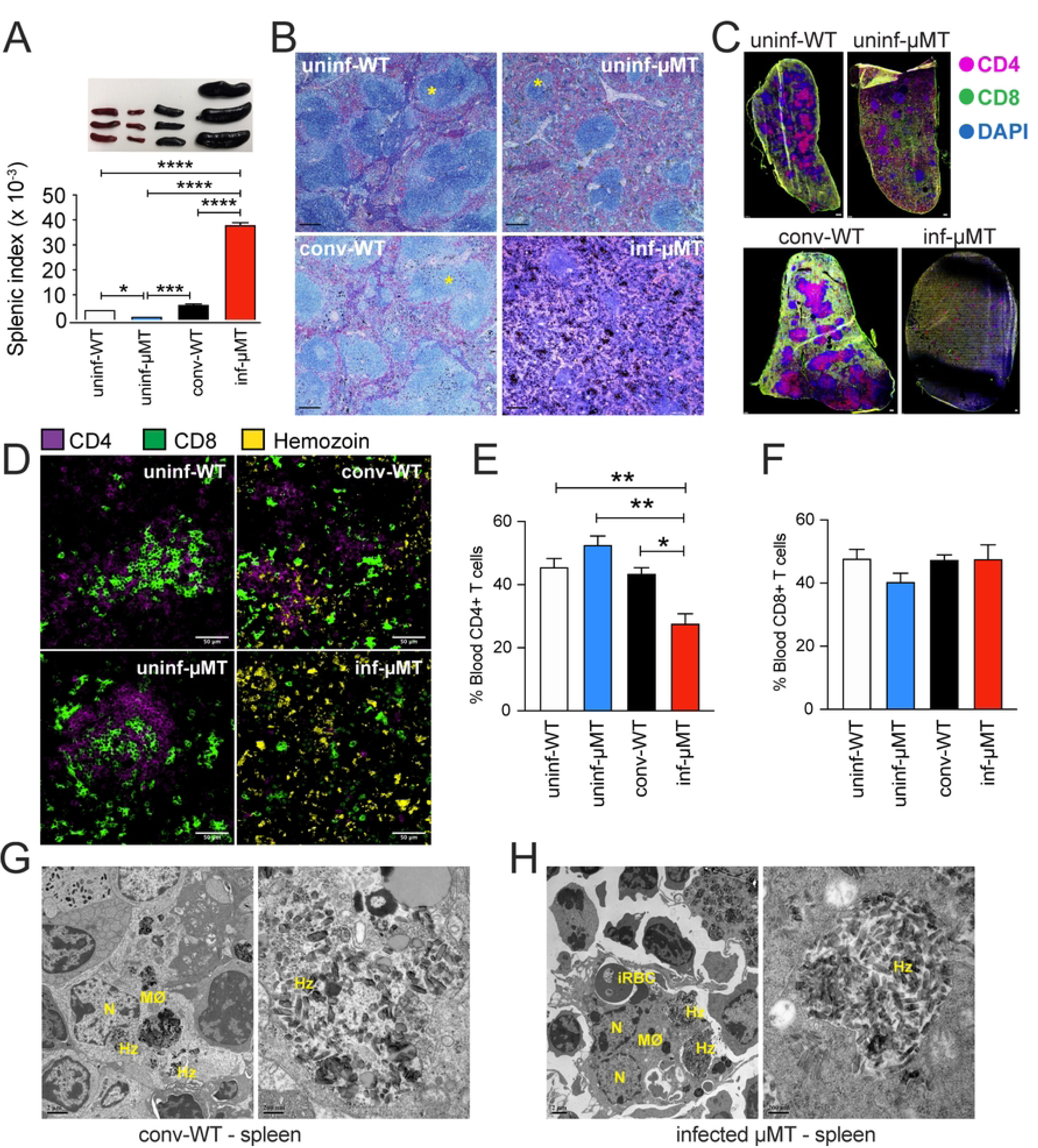
Chronic infection induces dramatic changes in spleen microarchitecture and hemozoin accumulation in the spleen. (A) Representative images of spleens (upper) and the associated splenic index (spleen weight/body weight) (bottom) from uninfected WT (uninf-WT, white bar), uninfected μMT^−/−^ (uninf-μMT^−/−^, blue bar), infected WT (inf-WT [convalescent], black bar), and *P. chabaudi*-infected μMT^−/−^ (inf-μMT^−/−^, red bar) mice 99 days post-infection (dpi), n=3. (B) Paraffin-embedded spleen sections H&E stained from uninf-WT, uninf-μMT^−/−^, inf-WT (convalescent), and inf-μMT^−/−^ mice 99 dpi. White pulps are indicated (*). Bar = 200 μm. (C) Fluorescence image from splenic sections (tile scan mode) stained with anti-CD4 (magenta), anti-CD8 (green), and DAPI (blue) from uninfected WT, uninfected μMT^−/−^, infected WT (convalescent), and infected μMT^−/−^ mice 99 dpi, bar = 500 μm. (D) Confocal microscopy images from splenic sections stained with anti-CD4 (magenta), anti-CD8 (green), and hemozoin (yellow) were visualized by confocal reflection microscopy from uninf-WT, uninf-μMT^−/−^, inf-WT (convalescent), and inf-μMT^−/−^ mice 99 dpi, bar = 50 μm. (E) Percentage of CD4+ and (F) CD8+ cells from peripheral blood of uninf-WT (white bar), uninf-μMT^−/−^ (blue bar), inf-WT (convalescent) (black bar), and inf-μMT^−/−^ (red bar) mice 99 dpi, n=3. (G) Transmission electron microscopy of spleen sections from inf-WT (convalescent) mice showing a macrophage (MØ), nucleus (N), and hemozoin crystals (Hz); bar = 2 μm (left panel). A high magnification of hemozoin crystals is shown in the right panel (bar = 200 nm). (H) Transmission electron microscopy of spleen sections from infected μMT^−/−^ mice showing a macrophage (MØ), infected red blood cell (iRBC), nuclei (N), hemozoin crystals (Hz), bar = 2 μm (left panel), and a high magnification of hemozoin crystals, bar = 200 nm (right panel). Data in A, E, and F are shown as mean ± SEM with * P<0.05, ** P<0.01, *** P<0.001, and **** P<0.0001, as determined by one-way ANOVA followed by Tukey’s multiple comparisons test.

As *Plasmodium* spp. infection temporarily changes the distribution of splenic cells during acute infection, we evaluated the distribution of CD4+ and CD8+ cells in splenic tissue sections by fluorescence microscopy at 99 days p.i. We observed that uninfected μMT^−/−^ mice had an altered distribution of T cells, as well as a decrease in the frequency of CD4+ and CD8+ T cells compared to uninfected WT mice (Fig 3C). Convalescent WT mice infected with *P. chabaudi* 99 days prior preserved the same frequency and distribution of CD4+ and CD8+ cells in their spleens (Fig 3C). In contrast, chronically infected μMT^−/−^ mice had a dramatic loss of standard splenic microarchitecture (Fig 3C), characterized by an irregular distribution of CD4+ and CD8+ T cells (Fig 3D). In addition, a reduction in blood CD4+ T cells, but not CD8+ T cells, was observed in chronically infected μMT^−/−^ mice (Figs 3E and 3F). Anti-CD11c and anti-F4/80 staining of tissue sections showed that chronically infected μMT^−/−^ mice had reduced percentages of dendritic cells (DCs) and increased macrophages as well as massive hemozoin accumulation (Fig. 3D and S4 Fig). The presence of large quantities of hemozoin is quite interesting, given that many groups have observed that hemozoin is potently proinflammatory (Arese and Schwarzer, 1997; Coban et al., 2005; Parroche et al., 2007) and appears to be biologically active for months post-infection (Frita et al., 2012).

To better visualize the structures in which hemozoin crystals accumulate in the spleens of chronically infected μMT^−/−^ mice, sections of this organ were processed and analyzed by transmission electron microscopy (TEM). We examined the accumulation of hemozoin in the spleens of WT and μMT^−/−^ mice at 99 days p.i. and observed regions of dark pigment in splenic sections in both chronically infected μMT^−/−^ and convalescent WT mice (Fig 3G), indicating ongoing hemozoin accumulation. Although seen less frequently, phagocytes with accumulated hemozoin crystals were observed in convalescent WT mice spleens (Fig 3G). In contrast, chronically infected μMT^−/−^ mice frequently showed phagocytosed hemozoin crystals and infected RBCs inside phagocytes (Fig 3H, left panel). High magnification showed large accumulations of hemozoin crystals inside giant phagocytes in splenic sections from chronically infected μMT^−/−^ mice (Fig 3H, right panel). Analysis of confocal images in a Z-plane from spleen sections of chronically infected μMT^−/−^ mice showed that hemozoin crystals colocalized within both DCs and macrophages (S4 Fig, right panel). Hence, long-term infection with *Plasmodium* resulted in the massive accumulation of an immunologically active crystal without symptomatology.

### Chronically infected μMT^−/−^ mice have persistently high plasma levels of IL-10, TNF-α, and IFN-γ

During the acute phase of *P. chabaudi,* proinflammatory cytokines are produced, induced by damage-associated molecular patterns (DAMPs) and pathogen-associated molecular patterns (PAMPs) released during the *Plasmodium* spp. erythrocytic cycle. To assess if cytokines were released during chronic *P. chabaudi* infection, plasma samples from chronically infected μMT^−/−^ mice, convalescent WT, uninfected WT, and μMT^−/−^ control groups were assessed by a bead-based multiplex immunoassay on day 99 p.i. Chronically infected μMT^−/−^ mice had increased levels of IL-10, TNF-α, and IFN-γ (Figs 4A-C). No differences in G-CSF and IL-6 circulating levels were observed amongst the groups (Figs 4D-E). Although CD4+ T cells are considered to be the major producer of IL-10 in acute *Plasmodium* infection, we observed no difference in IL-10 production amongst CD4+, CD4-, or CD14+ cells in mice peripheral blood (S5 Fig). We assessed IL-10 levels in the total spleen lysates and observed an increase in IL-10 expression in chronically infected μMT^−/−^ mice compared with all the other groups (S6 Fig). Thus, despite the apparent asymptomatic nature of the infection, chronically infected μMT^−/−^ mice showed persistent production of the anti-inflammatory cytokine IL-10 as well as proinflammatory cytokines.

**Fig 4.**
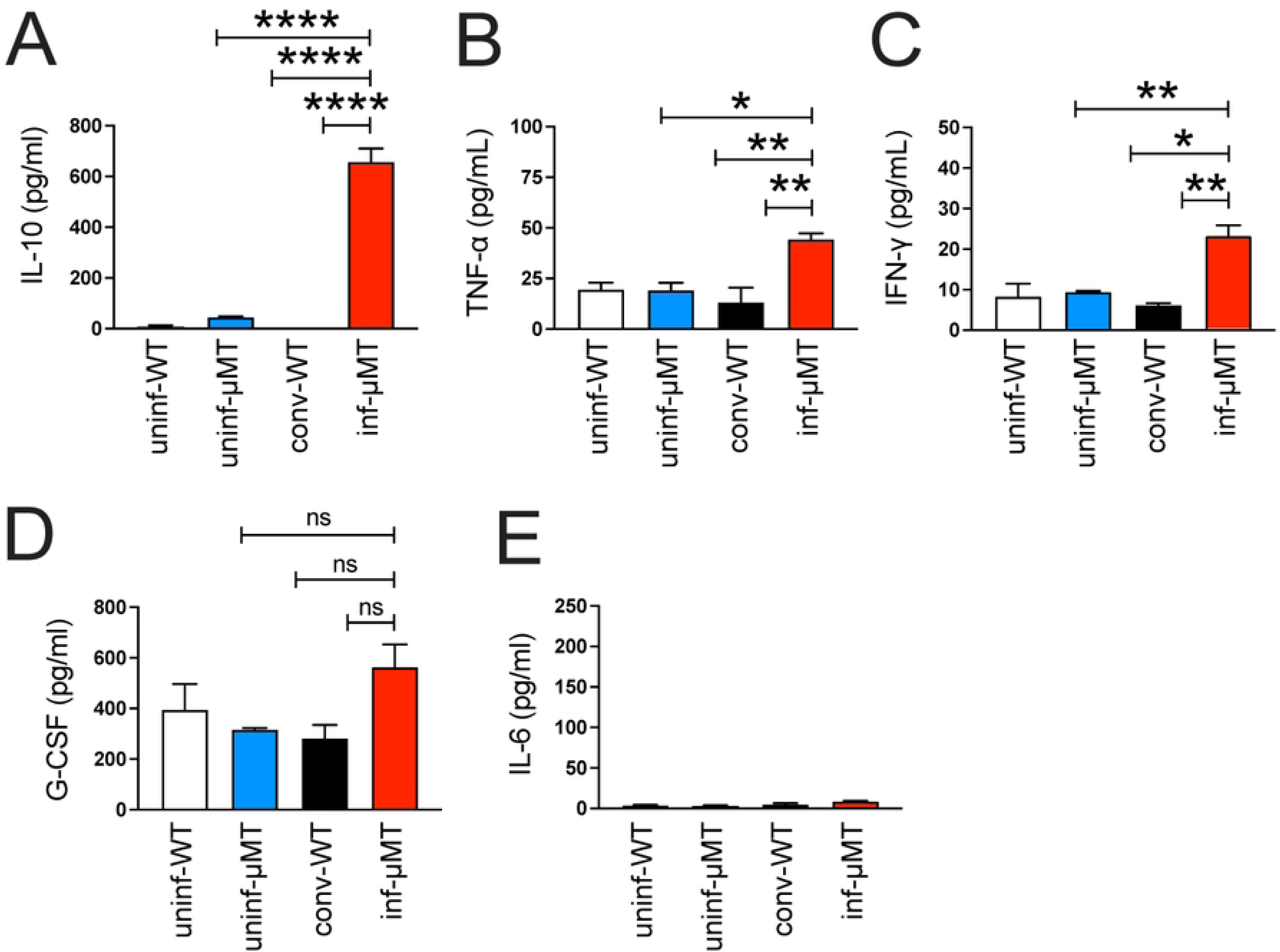
Chronically infected µMT^−/−^ mice have high plasma levels of IL-10, TNF-α, IFN-γ, and G-CSF. Plasma levels of (A) IL-10, (B) TNF-α, (C), IFN-γ, (D) G-CSF, and (E) IL-6 from uninfected WT (uninf-WT, white bars), uninfected μMT^−/−^ (uninf-μMT^−/−^, blue bar), convalescent infected WT (conv-WT, black bar), and *P. chabaudi*-infected μMT^−/−^ mice (inf-μMT^−/−^, red bar) 99 days p.i. (dpi) were assessed by bead-based multiplex immunoassay, n=3. Data are shown as mean ± SEM. * P<0.05, ** P<0.01, *** P<0.001, **** P<0.0001, as determined by one-way ANOVA followed by Tukey’s multiple comparisons test.

### IL-10 protects μMT^−/−^ mice chronically infected with *Plasmodium chabaudi* by preventing excessive inflammatory responses

Studies in mice and humans demonstrate that enhanced production of IL-10 typically accompanies acute *Plasmodium* infection (Freitas do Rosario et al., 2012; Turner et al., 2021), suggesting that this cytokine modulates the degree of inflammation. To test the hypothesis that the high plasma levels of IL-10 observed in chronically infected μMT^−/−^ mice protect against observable disease, chronically infected μMT^−/−^ mice received injections of anti-IL-10R or an isotype control antibody (S7 Fig). Blocking IL-10 signaling rapidly reduced the parasitemia level after the first anti-IL-10R antibody injection, and it remained low for nearly 2 weeks (Fig 5A). Moreover, a significant loss of body weight was observed after two weeks of treatment (Fig 5B). In addition, IL-10R blockade increased the numbers of monocytes and neutrophils in the peripheral blood ∼2 weeks post-antibody injection (Figs 5C–5D). The presence of an overall inflammatory state was confirmed by the levels of C-reactive protein that started to increase 7 days post-antibody injection (Fig 5E). After day 16 post-antibody injection, infected μMT^−/−^ mice that received anti-IL-10R injections began to die (Fig 5F). Fluorescence images from whole spleen tissue sections stained with anti-Ly-6C/Ly-6G showed increased polymorphonuclear neutrophil (PMN) infiltration in the spleen 21 days post-antibody injection (Fig 5G). Altogether, these results demonstrated that blocking IL-10 production in chronically infected μMT^−/−^ mice reduced parasitemia but resulted in a lethal excessive proinflammatory response.

**Fig 5.**
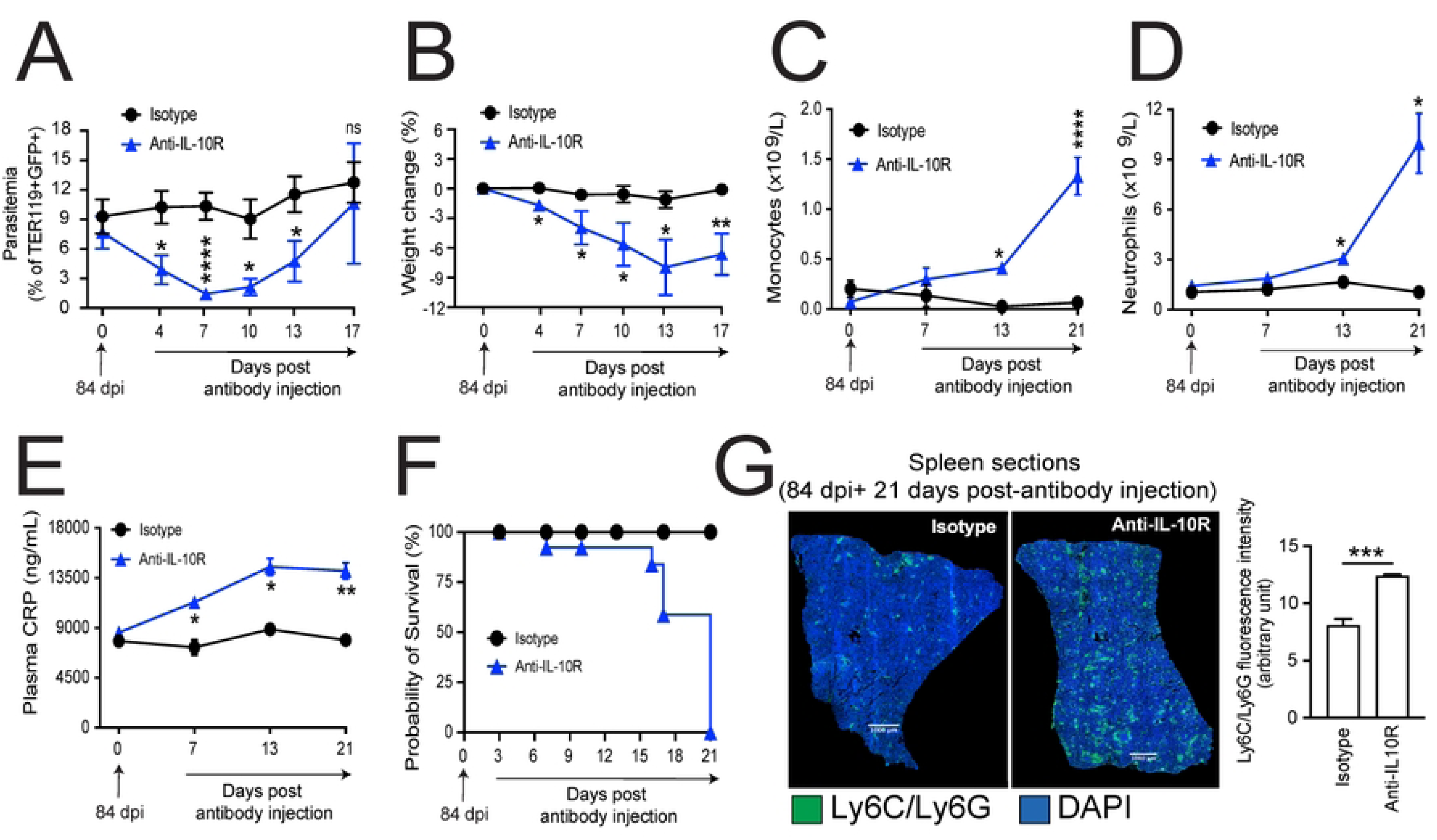
IL-10 protects μMT^−/−^ mice chronically infected with *Plasmodium chabaudi* by preventing excessive inflammatory responses. Chronically infected μMT^−/−^ mice were treated via intraperitoneal injection with an anti-IL-10 receptor mAb or an isotype Ab control 84 d.p.i. Data were recorded before Ab (anti-IL-10R or isotype control) administration (day 0, at 84 d.p.i.) and 4, 7, 10, 13, and 17 days after. (A) Parasitemia is shown as a percentage of TER119^+^GFP^+^ events, and (B) body weight is shown as percentage change. (C) Number of monocytes (n=3), and (D) neutrophils (n=3) were assessed by complete blood count (CBC) from blood samples from infected μMT^−/−^ mice before Ab (anti-IL-10R or isotype control) administration (day 0, at 84 d.p.i.) and 7, 13, and 21 days after Ab injection. (E) Plasma C-reactive protein levels in ng/mL (n=4) were assessed before Ab (anti-IL-10R or isotype control) administration (day 0, at 84 d.p.i.) and 7, 13 and 17 days after Ab (n=4). (F) Kaplan Meier survival curve of treated mice (n=12). (G) Fluorescent microscopy image of whole spleen sections (tile scan mode) stained with DAPI (blue) anti-Ly6C/Ly6G antibody (green) 21 days after isotype (left panel) or anti-IL-10R (right panel) antibody injection, bar = 1000 μm. For (A–E), * P<0.05, ** P<0.01, *** P<0.001, **** P<0.0001, as determined by two-way ANOVA followed by Tukey’s multiple comparisons test (comparing isotype vs. anti-IL-10R), or by unpaired Student’s t-test (G graph bar).

### IL-10R blockade increased the plasma levels of proinflammatory cytokines

Antibody-mediated IL-10R blockage increased the plasma levels of IL-10 (Fig 6A), proinflammatory cytokines, TNF-α, IFN-γ, IL-6, and the chemokine G-CSF (Figs 6B–6E). Hence, the high circulating levels of IL-10 present in chronically infected mice that engage the IL-10R were responsible for suppressing several cytokines generally considered to be pro-inflammatory in a manner that suggests a regulatory feedback loop.

**Fig 6.**
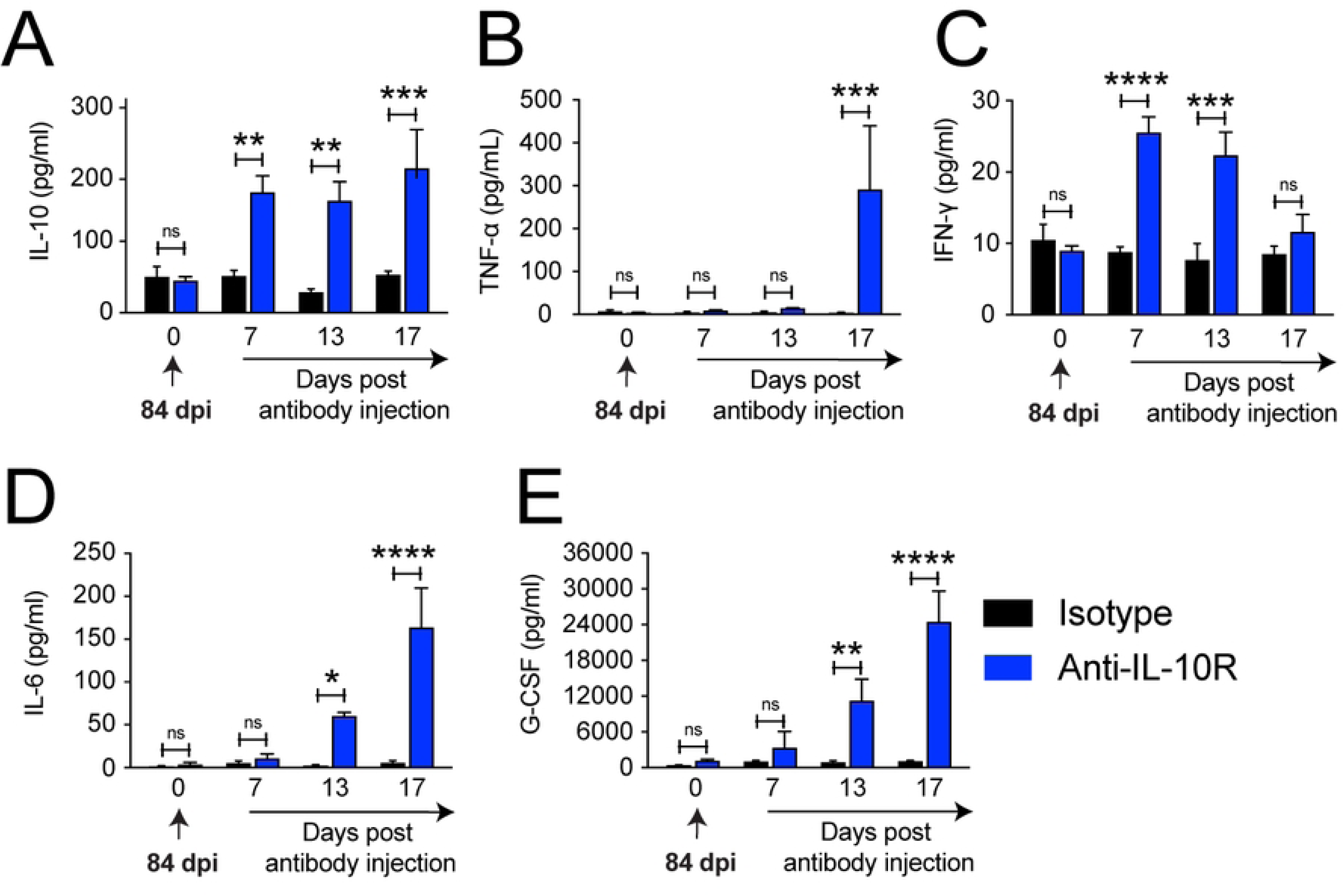
IL-10R blockage increases the plasma levels of IL-10, TNF-α, IFN-γ, IL-6, and G-CSF. Circulating levels in pg/mL of (A) IL-10, (B) TNF-α, (C) IFN-γ, (D) IL-6, and (E) G-CSF were assessed before (day 0) and at days 7, 13, and 17 after isotype (black bars, n=4) or anti-IL-10R (blue bars, n=4 or at day 17 n=3) antibodies were injected into chronically infected μMT^−/−^ mice (84 dpi, at day zero). Data are shown as mean ± SEM with * P<0.05, ** P<0.01, *** P<0.001, and **** P<0.0001, as determined by a two-tailed, unpaired Student’s t-test.

### Bone marrow macrophages from chronically infected μMT^−/−^ mice have differential chromatin accessibility

To assess if chronic *Plasmodium* spp. infection induces changes in chromatin accessibility in phagocytes, bone marrow macrophages from convalescent WT, chronically infected μMT^−/−^ mice 117 days p.i., and uninfected control groups were isolated to a high degree of purity by FACS and assessed by Assay for Transposase-Accessible Chromatin using sequencing (ATAC-seq). We chose to study bone marrow macrophages because of their principal role in inflammation during malaria, the observation that they harbored abundant amounts of parasite and hemozoin (see S4 Fig), their relative abundance, and the ease by which they can be purified. Principal component analysis of ATAC-seq consensus peaks showed that infected μMT^−/−^ mice possessed a distinct accessibility profile from the other groups (Fig 7A). Consensus peak annotation of each genomic region showed more than 25% of the accessible regions were promoters (Fig 7B). Indeed, despite the lack of symptomatology in chronically infected μMT^−/−^ mice, innate immune response promoters and promoter regions involved in cytokine production were clearly areas of enhanced chromatin accessibility, as demonstrated by gene ontology (GO) term analysis (Fig 7C). Normalized counts for promoter regions in selected immune response genes displayed significant differences in chromatin accessibility for genes, such as *Ccl12*, *Map4k4*, *Il27*, and *Stat3*, although we did not observe differences in chromatin accessibility in the promoters of the *Il10* or *Il10* receptor genes (Fig 7D).

**Fig 7.**
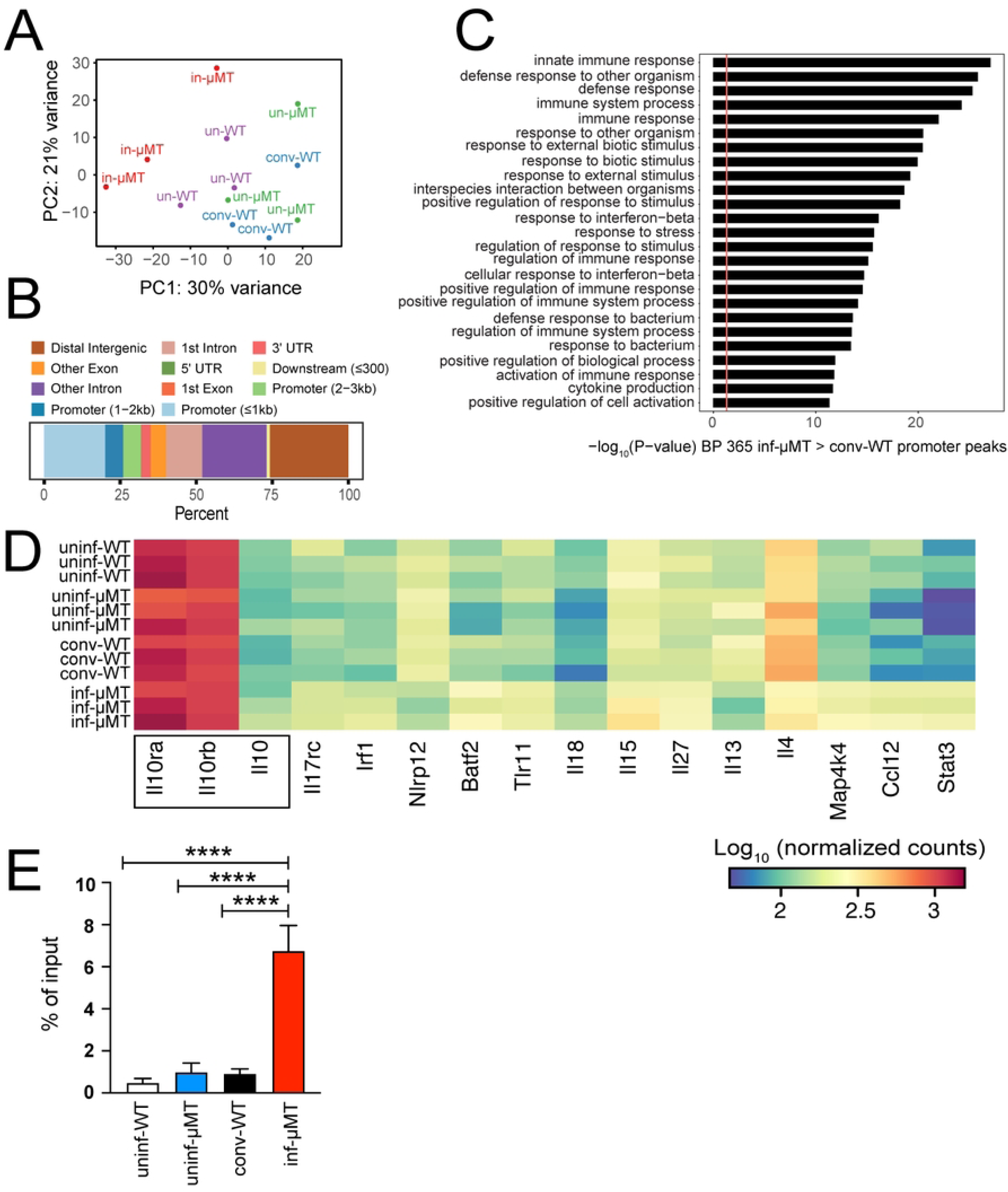
Bone marrow macrophages from chronically infected μMT^−/−^ mice have differentially accessible chromatin. (A) Principal component analysis plot of ATAC-seq consensus peaks (regularized log-transformed counts) from bone marrow-derived macrophages from uninfected WT (un-WT, magenta), uninfected μMT^−/−^ (un-μMT, green), convalescent WT (conv-WT, blue), and infected μMT^−/−^ (in-μMT, red) mice 117 days p.i. (dpi). Each point represents an independent biological replicate. (B) Consensus peak annotation showing the percentage of peaks for each genomic region. (C) Significantly enriched GO terms for genes that had a promoter peak (within -1000 bp and +500 bp from a transcription start site) that was significantly greater in *P. chabaudi*-infected μMT^−/−^ than convalescent WT mice (adjusted P-value < 0.05, n=365). (D) Heat map of normalized counts for promoter regions in selected immune response genes with significant differences in chromatin accessibility between *P. chabaudi*-infected μMT^−/−^ and convalescent WT mice (adjusted P-value < 0.05). The peaks are ordered by adjusted P-value with the most significant peaks on the right. The *Il10* genes are boxed (P-values>0.05) and included for comparison for uninf-WT, uninf-μMT^−/−^, conv-WT, and inf-μMT^−/−^ mice (3 mice/group). (E) qPCR for the *Il10* promoter from chromatin immunoprecipitated with anti-mouse STAT3 monoclonal antibody of bone marrow macrophages from uninf-WT (n=11), uninf-μMT^−/−^ (n=11), conv-WT (n=10), and inf-μMT^−/−^ (n=12) mice. Data are shown as mean ± SEM with **** P<0.0001, as determined by one-way ANOVA followed by Tukey’s multiple comparisons test.

Because STAT3 regulates IL-10 production in macrophages (Lucas et al., 2005; Staples et al., 2007), we evaluated whether the increased accessibility of the *Stat3* promoter might lead to an increase in STAT3 binding to the *Il10* promoter of bone marrow macrophages. We performed a chromatin immunoprecipitation (ChIP) assay using monoclonal anti-mouse STAT3 antibody followed by qPCR using primers targeting the *Il10* promoter. On average, more than a five-fold increase of STAT3 binding to the *Il10* promoter was observed in chronically infected µMT^−/−^ mice compared to any of the other groups of mice (range of 2–14-fold, \*\*\*\**P*<0.0001; Fig 7E).

## Discussion

Subclinical malaria is the most common condition in malaria-endemic areas of the world. Often referred to as “asymptomatic” malaria, subclinical malaria has been hotly debated (Andolina et al., 2021). Many experts feel that it is not truly asymptomatic, but rather that its symptoms are often challenging to identify (Chen et al., 2016). Patients with “asymptomatic” malaria might suffer from chronic anemia, nephrotic syndrome, co-infection with a potentially lethal bacterium, cognitive impairment, and other problems. Conversely, subclinical malaria protects children against cerebral malaria, life-threatening anemia, and other life-threatening manifestations of the disease (Akiyama et al., 2016). Many physicians propose that subclinical disease confers immunity to severe disease, as the incidence of febrile illness over time may be reduced in populations of infected asymptomatic individuals (Buchwald et al., 2019; Males et al., 2008; Sonden et al., 2015). Thus, children with chronic *Pf* malaria are typically not treated because, by clinical measures, they appear normal; however, the prevalence of subclinical malaria poses a significant challenge to eradicating malaria (Cheaveau et al., 2019).

In this study, we investigated the immune regulatory mechanisms behind long-term *P. chabaudi* infection using the µMT^−/−^ mouse model that exhibits many characteristics of subclinical malaria in humans. First and foremost, after an acute infection, persistent patent parasitemia occurs for months in the absence of any signs of disease. Indeed, we report here that (after a brief period of initial distress), *P. chabaudi*-infected µMT^−/−^ mice have patent parasitemia for months. An inspection of the mice for any external signs of disease suggested that they were entirely asymptomatic for at least 7 months, at which point the degree of anemia becomes life-threatening. Our observations were confirmed by metabolic studies that revealed that the mice had no detectable alterations in physical activity or food and water intake after an initial decrease at day 8 p.i. These findings resemble, to some degree, subclinical disease in children with chronic *Pf* malaria. It is noteworthy that the levels of parasitemia observed in our study in asymptomatic chronically infected µMT^−/−^ mice (>10%) are higher than those observed in subclinical *Plasmodia* spp. infection in humans. However, elevated levels of parasitemia in chronic asymptomatic infection have been demonstrated before in a mouse model for chronic long-term infections using immunocompromised mice (Fontana et al., 2016).

Although chronically infected µMT^−/−^ mice have no apparent outward signs of disease, they exhibited ongoing hemolytic anemia, accompanied by profound reticulocytosis and an enormous deposition of hemozoin in the spleen, liver, and bone marrow. We also showed that these mice have dramatic alterations in the architecture of their spleen and massive splenomegaly. Recently, it was demonstrated that a large biomass of nonphagocytosed, intact RBCs infected with *P. falciparum* or *P. vivax* in all stages of development accumulate in the spleen of asymptomatic individuals at levels that are hundreds to thousands of times higher than the biomass of infected RBCs in the peripheral blood (Kho et al., 2021). This observation is consistent with the atypical splenomegaly observed in our chronically infected µMT^−/−^ mice that also showed all stages of nonphagocytosed, intact infected RBCs in the spleen tissue. In humans, a phenomenon known as hyperreactive malarial splenomegaly syndrome (HMS) is induced by chronic stimulation of *Plasmodium* spp. antigens (Leoni et al., 2015; McGregor et al., 2015). Moreover, high circulating levels of IL-10 are a feature of malaria patients with HMS (Alkadarou et al., 2013). The μMT^−/−^ mouse model of asymptomatic malaria also mirrored the correlation between IL-10 levels and severity observed in other studies of *Plasmodium* infections in both humans (Boyle et al., 2017; Wilson et al., 2010) and mice (Li et al., 1999). In addition, B-cell-deficient (J_H_^−/−^) mice chronically infected by *P. chabaudi adami* (Weidanz et al., 2005) have high circulating levels of IL-10. The main source of IL-10 in long-term malaria is still unclear. Although previous studies have shown that specific populations of T cells are the source of IL-10 (Couper et al., 2008; Freitas do Rosario et al., 2012), our results showed no difference in IL-10 production from CD4+ T cells from peripheral blood. The increased IL-10 production seems to take place in the spleen, where we observed higher production of IL-10 in the tissue lysate.

IL-10 is produced by the vast majority of immune cells, both from the innate immune system and from the adaptive immune system (Kumar et al., 2019), including peripheral blood monocytes and CD4+ T cells. During *P. falciparum* infection, these cells continue to produce cytokines in response to malarial antigens for prolonged periods, even in the absence of parasite transmission (Moormann et al., 2006; Moormann et al., 2009; Wipasa et al., 2011). Previous reports suggest there is a burst of IL-10 production by DCs and macrophages after the phagocytosis of natural hemozoin (Bobade et al., 2019). We observed colocalization of DCs and macrophages with hemozoin crystals in the spleen of chronically infected μMT^−/−^ mice, suggesting that these cells could be the source of high circulating levels of IL-10. In children, uncomplicated acute *P*. *falciparum* infection is related to CD4 T cell production of IL-10 without the production of the pro-inflammatory cytokines IFN-γ and TNFα (Boyle et al., 2017). During acute infection, IFN-γ+ Th1 cells seem to be the primary producers of IL-10. Although we cannot rule out the production of IL-10 by IFN-γ+ Th1 cells, these cells were not seen in abundance when blood samples from chronically infected µMT^−/−^ mice were examined by flow cytometry (S5 Fig). Thus, the primary source of IL-10 in chronically infected µMT^−/−^ mice is likely from a solid organ, such as the spleen and the liver. Indeed, we observed higher expression of IL-10 mRNA from chronically infected µMT^−/−^ mice in the liver (but not the spleen) (S6A and S6B Figs). Although Kupffer cells represent 35% of nonparenchymal cells in the liver of adult mice and account for 80–90% of resident macrophages (Gregory and Wing, 2002), the role of hepatocytes as a major source of IL-10 cannot be ruled out. Indeed, hepatocyte-derived IL-10 supports chronic murine trypanosomiasis in a manner that is highly reminiscent of *P. chabaudi* infection in µMT^−/−^ mice (Stijlemans et al., 2020). At this point, it is impossible to confidently say what cell type is the main contributor to the production of IL-10 in chronically infected asymptomatic µMT^−/−^ mice. In addition, while it is widely believed that T lymphocytes are the most important source of IL-10 in humans with subclinical *P. falciparum*, the role of solid organs in maintaining chronic parasitemia is clearly worth examining.

One might hypothesize that the chronically infected phenotype would be accompanied by the silencing of the innate immune system, perhaps by inactivating TLRs or other pattern recognition receptors involved in immunity to *Plasmodium*. However, our data show that the innate immune response is never completely silenced in chronically infected animals. Indeed, many proinflammatory cytokines are elevated in an ongoing manner throughout the course of the disease. One might also expect parasitemic mice to be compromised by elevated levels of IFN-γ, IL-12, and other cytokines. However, as is often the case during inflammation, IL-10 was persistently elevated in these mice, resulting in a state of resistance to disease. Research suggests that IL-10 protects the host against excessive inflammation but may impede parasite clearance, possibly by inhibiting dendritic cell and macrophage functions (O’Farrell et al., 1998). IL-10-deficient mice infected with a non-lethal *P*. *chabaudi* AS strain showed exacerbated pathology with excessive proinflammatory responses and developed cerebral edema and hemorrhages (Li et al., 1999; Sanni et al., 2004). We demonstrate that IL-10 enables low levels of parasitemia to persist in asymptomatic μMT^−/−^ mice and that blocking IL-10 transiently decreased the infection but results in an excess inflammatory response that is lethal.

*Plasmodium falciparum* manipulates the host’s immune system genes through histone modifications and chromatin remodeling (Schrum et al., 2018). Our lab and others previously demonstrated that *Plasmodium* spp. infection induces epigenetic reprogramming in human monocytes and macrophages (Guha et al., 2021; Schrum et al., 2018). We now report our findings indicating that IL-10 production in chronic asymptomatic malaria is regulated by epigenetic changes. It is known that IL-10 can induce IL-10 production by macrophages through a Stat3-dependent mechanism (Staples et al., 2007), and this result is consistent with our ATAC-seq data that showed increased accessibility of the *Stat3* promoter region in chronically infected µMT^−/−^ mouse macrophages. In addition, chronically infected µMT^−/−^ macrophages had increased STAT3 binding to the *Il10* promoter region. However, whether histone modifications alter the accessibility of the promoter regions of *STAT3* in chronically infected mice remains unknown. An equally important question is whether the upstream events leading to the epigenetic changes that sustain high levels of IL-10 are due to the same signals (for example, the activation of a TLR) that result in the production of proinflammatory cytokines in acute disease.

Hutchins and colleagues demonstrated that IL-10 stimulation of macrophages activates the IL-10R/STAT3 signaling pathway to drive an anti-inflammatory response by inducing genes that repress the transcription of proinflammatory cytokine genes (Hutchins et al., 2012). They also showed that the IL-10R/STAT3 axis negatively regulates macrophage responses (Hutchins et al., 2012). That the apparently asymptomatic state in the µMT^−/−^ mice was due to the effects of IL-10 on the phagocytic compartment is strengthened by our observation that IL-10R blockade increases circulating monocytes and neutrophils that presumably account for the concordant decrease in parasitemia. Future studies are needed to examine how IL-10 and proinflammatory cytokines are regulated during asymptomatic malaria to identify strategies to clear the parasite while simultaneously preventing the deleterious excessive immune response.

In summary, we demonstrated that chronic *P. chabaudi* infection induced substantial accumulation of hemozoin in the spleen, splenomegaly, remodeling of splenic architecture cell distribution, and changes in leukocyte frequency. Despite an apparent lack of symptoms, infected animals had high circulating levels of inflammatory mediators and IL-10, apparently maintained by epigenetic changes. These high levels of IL-10 protected mice from a severe inflammatory response despite the failure to control the parasite burden. Our data, combined with observations of malaria disease in humans, suggest a real relevance to the results we report here in a chronically infected mouse model, in which we observed persistently elevated levels of IL-10 that prevent death due to *Plasmodium* parasitemia, albeit at an obvious cost in terms of hypersplenism and severe anemia. The epigenetic findings in chronically infected mice suggest a mechanism that can be tested in humans—that enhanced production of Stat3 drives IL-10 production via transcriptional activation (Benkhart et al., 2000). Finally, both IL-10 and IL-27 are anti-inflammatory, and their expression needs to be examined as a cause for the increased susceptibility to bacterial infections, such as *Salmonella* spp., that has been demonstrated in children with asymptomatic *P. falciparum* infection (Post et al., 2021).

## Materials and methods

### Ethics statement

All animal experiments were conducted in accordance with the guidelines of the American Association for Laboratory Animal Science (AALAS) and approved by the Institutional Animal Care and Use Committees (IACUC) at the University of Massachusetts Chan Medical School under protocol number 202000093.

### Mice and *Plasmodium* infection

Homozygous mutant *Ighm^tm1Cgn^ mice*, also known as µMT^−/−^ mice (lacking mature B cells due to a disruption of the μ gene) and C57BL/6J WT control mice, ages 8–12 weeks, were obtained from The Jackson Laboratory and housed in ventilated cages with standard food and water *ad libitum* and at 12 hours light/dark cycle. Male and female mice were used throughout the experiments.

The *Plasmodium chabaudi* AS strain expressing the green fluorescent protein (GFP) (Reece and Thompson, 2008) was obtained from Dr. Joanne Thompson from the University of Edinburgh, UK and the European Malaria Reagent Repository (www.malariaresearch.eu). A vial from deep-frozen stock was used to start the primary infection in C57BL/6J mice. The parasite was maintained *in vivo* by weekly passages for a maximum of ten weeks. After 2 or 3 passages, a suspension of 10^5^ infected RBCs in 1XPBS (maximum volume of 200 µl) was injected intraperitoneally (i.p.) in µMT^−/−^ and C57BL/6J mice. Parasitemia was monitored weekly from 1.5 µl of tail blood samples by flow cytometry using the gating strategy represented in S2A Fig.

### Flow cytometry

To evaluate parasitemia and/or reticulocytes, 1.5 µL of whole blood was extracted from tail bleeds, diluted in 1XPBS + 1% fetal bovine serum (FBS), and fixed with 0.025% glutaraldehyde in 1XPBS for 20 minutes. After centrifugation at 300*g* for 5 min at 4 °C, the pellet was resuspended in staining solution and kept on ice for 35 minutes, protected from the light. The cells were washed twice in 1XPBS + 1% FBS and resuspended in 200 µl of 1XPBS + 1% FBS. Data were acquired on an LSRII cytometer (BD Biosciences) using DIVA software (BD Biosciences) and analyzed by FlowJo software version 10 (FlowJo LLC, Ashland, OR).

### Intracellular staining

Intracellular cytokines were detected after stimulation with 20 ng/mL phorbol myristate acetate (PMA), 1 μg/mL ionomycin, and 5 μg/mL brefeldin A for 6h. Surface labeling with anti-CD4-Pacific blue and anti-CD14-BV421, as well as viability dye Zombie NIR, were performed before cell fixation with paraformaldehyde (BioLegend #424401), followed by permeabilization buffer (BioLegend #424401). To detect intracellular cytokines, the cells were further stained with anti-IL-10–PE (BioLegend JES5-16E3).

### Nested polymerase chain reaction

A nested PCR targeting *Plasmodium* spp. chromosome 12 was developed to detect sub-patent parasitemia. Serial dilutions (from 1:10 to 1:10^6^) of whole blood samples from infected mice (0.1% of parasitemia) were used to test the limit of detection. A calibration with known parasite densities was used as a reference. The detection limit was roughly ≥ 1 parasite/μL at the dilution 1:10^4^ (S2B Fig).

DNA amplification was performed using a Phusion Blood Direct PCR Kit (Thermo Fisher Scientific, #F547F), which does not require DNA purification. Two sets of primers targeting the *Plasmodium* spp. chromosome 12 were used. The outer primers at 0.25 µM were: Forward: (OPFWD) 5’-GCCTTCCTTAGATGTGGTAGC-3’ and Reverse: (OPRVS) 5’-TGATCTTGCCAGTAGTCATATGC-3,’ and the inner primers at 0.50 µM were: Forward: (IPFWD) 5’-CGTTACCCGTCATAGCCATG-3’ and Reverse: (IPRVS) 5’-CGAACGGCTCATTAAAACAGT-3’ (see Key resources table). For the outer PCR reaction, 4 µl of whole blood obtained from tail bleeds were used in a 20 µl final volume reaction. For the inner PCR, 1 µl of the amplicon generated by the outer PCR product was used as a sample in a 20 µl of final volume reaction. The following conditions were used: outer PCR: 98 °C/5 min, [98 °C/1 sec, 62 °C/5 sec, 72 °C/15 sec] 25X; 72 °C/1 min; 4 °C hold. Inner PCR: 98 °C/5 min, [98 °C/1 sec, 63 °C/5 sec, 72 °C/15 sec] 30X, 72°C/1 min, 4 °C hold. The size of the PCR product was predicted as 274 bp and confirmed in a 2% agarose gel in 1x Tris Acetate-EDTA (TAE) buffer (Sigma, T9650), stained by ethidium bromide (MP Biomedicals, LLC – Cat. 802511), and a 100 bp DNA ladder (Thermo Scientific SM0243) was used as a reference. An agarose gel loading dye 6X, glycerol-based (Boston Bioproducts – Cat. BM-100G), was added to samples which ran for 45 minutes at 124V in an Eppendorf thermocycler Mastercycler nexus GX2, and the images were acquired in a ChemiDoc Touch Imaging System (Bio-Rad).

### Food and water intake, energy expenditure, and physical activity

Mice were housed individually in metabolic cages (TSE Systems, Chesterfield, MO) at the National Mouse Metabolic Phenotyping Center (MMPC) at UMass Chan Medical School. Food and water intake, spontaneous physical activity, and indirect calorimetry were simultaneously measured in awake mice. Indirect calorimetry was assessed by evaluating the production of carbon dioxide and oxygen consumption, and (based on these values) the respiratory exchange ratio and energy expenditure were calculated. Physical activity was assessed by measuring the horizontal and vertical movements of the mice using a three-dimensional series of infrared beams (X + Y + Z axis). Data from 3 consecutive days were recorded and were expressed as an average of 24 hours. The days chosen for analysis were based on the *P. chabaudi* infection profile of C57BL/6J mice: days -3, -2 and -1 prior to infection (baseline, referred to as 1 day before infection), days 6, 7, and 8 p.i. (acute stage, referred to as 8 days p.i.); days 28, 29, and 30 p.i. (chronic stage with sub-patent parasitemia, referred to as 30 days p.i.) and days 55, 56 and 57 p.i. (clearance stage, when inf-WT mice were PCR negative).

### Complete blood counting

Complete blood counting was performed in 100 µl of whole blood collected in EDTA tubes by cheek bleeding. Samples were analyzed in a veterinary hematology analyzer VetScan HM5 at the UMass Chan Medical School Department of Animal Medicine.

### Glucose measurement

Blood glucose levels were assessed in blood from tail bleeds using the Nova Max® glucometer. All blood collection was performed during the first 3 hours of the mice light-cycle.

### Histology

Spleen and liver tissue sections (5 µm thick) were embedded in paraffin and stained with H&E by the Morphology Core at UMass Chan Medical School. For fluorescence microscopy, spleen and liver tissue sections (5 µm thick) were embedded in Tissue-Tek O.C.T. compound and fixed for 30 min in 4% paraformaldehyde, washed 3 times with 1XPBS, blocked for 1.5 hours in 1XPBS + 5% BSA + 1% serum from secondary antibody species, stained overnight at 40 °C protected from the light with primary antibodies, washed 3 times with 1XPBS, and stained for 1.5 hours at room temperature with secondary conjugated antibodies. Next, tissue sections were washed 3 times with 1XPBS, stained with DAPI for 5 min, washed 3 times with 1XPBS and covered with a thin layer of mounting compound (ProLong Gold Antifade Mountant, Thermo Fisher Scientific) and coverslips. The images were acquired using a Leica SP8 confocal laser scanning inverted microscope through confocal reflection and conventional fluorescence microscopy. Hemozoin images were acquired by reflection by placing the detector channel directly over the wavelength of the selected laser channel for reflection light capture, and the acusto-optical beamsplitter (AOBS) was set to allow 5–15% of laser light into the collection channel as previously described (Hornung et al., 2008).

### Transmission electron microscopy

Fresh spleen samples were fixed with a double-aldehyde fixative, 2.5% glutaraldehyde and 4% paraformaldehyde. Spleen samples were processed by the Electron Microscopy Core Facility at the UMass Chan Medical School. The images were acquired using a Philips CM10 transmission electron microscope (Field Electron and Ion Co., USA).

### Multiplex bead-based assays

Plasma from control and infected mice was used at a 1:2 and 1:4 dilution with the Bio-Plex Pro Mouse Cytokine 23 Plex Immunoassay (Bio-Rad Laboratories, Inc., CA, USA) following the manufacturer’s instructions. Standards provided with the kits as well as our samples were assayed in duplicate on a BioPlex 200 Luminex Instrument using the BioPlex manager software. A minimum of 50 beads of each analyte was acquired, and the median fluorescent intensity (MFI) was recorded. Quantitative values (pg/ml) were obtained using the 4-parameters standard curve of each cytokine/chemokine included in the kit. Results within the range of the standard were then used for statistical analysis.

### Detection of plasma cytokines and C-reactive protein

IL-27 plasma levels were assessed using the Mouse Cytokine 23-Plex immunoassay (Bio-Rad Laboratories, Inc., CA, USA) and by the IL-27 Mouse ELISA kit (Thermo Fisher Scientific, MA, USA) according to the manufacturer’s instructions. Plasma C-reactive protein levels were assessed by a Mouse C-Reactive Protein/CRP Quantikine ELISA Kit (R&D Systems, Inc., MN, USA) according to the manufacturer’s instructions.

### Quantitative real-time RT-PCR

Total RNA from the spleen or liver was extracted using the RNeasy Mini Kit (250) (Qiagen), and cDNA was synthesized using iScript reverse transcription Supermix for RT-PCR (Bio-Rad Laboratories, Inc., CA, USA). The qPCR reactions were performed using iTaq Universal SYBR Green Supermix (Bio-Rad Laboratories, Inc., CA, USA) and 250 nM of *Il10* forward and reverse primers (listed in the Key resources table). Mouse IL-10 mRNA relative expression was performed by the comparative threshold cycle (ΔC_T_) method and normalized using as a reference endogenous control primers (250 nM each, forward and reverse) targeting the mouse r18S gene.

### In vivo antibody treatment

Chronically infected μMT^−/−^ mice were separated into two groups: one group received monoclonal anti-mouse IL-10R antibody (clone 1B1.3A), and the second group received monoclonal isotype control, anti-horseradish peroxidase antibody (clone HRPN). The antibodies were delivered by intraperitoneal injection in 4 doses separated by 3 days. For the first two doses, 500 µg per mouse were administered (days 0 and 4); the subsequent two doses were 250 µg per mouse (day 7 and 10) (S7 Fig) as previously described (Gonzalez-Lombana et al., 2013).

### Bone marrow-derived macrophage isolation, transposition reaction and DNA purification

Cells from mice bone marrow were collected as previously described (Liu and Quan, 2015). Total mice bone marrow-derived cells Fc receptors were blocked with anti-CD16/CD32 antibody and then stained with anti-mouse CD11b eFluor 450, anti-mouse Ly6G-PerCP-Cy5.5, anti-mouse F4/80-APC, and Ghost viability dye BV510. Bone marrow macrophages were isolated by fluorescence-activated cell sorting (FACS) as CD11b+; Ly6G-; F4/80+ events. Then, 50,000 cells were centrifuged at 500*g* for 5 min at 4 °C. The pellet was washed once with 50 μl of cold 1XPBS and centrifuged at 500*g* for 5 min at 4 °C in a fixed-angle centrifuge. The supernatant was discarded, being careful to preserve the visible cell pellet by using two pipetting steps. The cell pellet was gently resuspended in 50 μl cold ATAC-Resuspension Buffer (RSB) containing three detergents (0.1% NP40, 0.1% Tween-20, and 0.01% digitonin) and homogenized by pipetting up and down 3 times and resting for 3 minutes, always on ice. Lysis buffer was removed by washing cells with 1 ml of cold ATAC-RSB containing only 0.1% Tween-20 (no NP-40 or digitonin) and inverting the tubes 3 times. Pelleted nuclei were obtained after centrifugation at 500*g* for 10 min at 4 °C. The supernatant was discarded, and the pellet was resuspended by pipetting up and down 6 times after adding 50 μl of a transposition mixture consisting of 25 μl 2x TD buffer (20 mM Tris-HCl pH 7.6, 10 mM MgCl_2_, 20% Dimethyl Formamide, water q.s.p.), 2.5 μl of a hyperactive form of Tn5 transposase loaded with Illumina adaptors (100 nM final), 16.5 μl PBS, 0.5 μl 1% digitonin, 0.5 μl 10% Tween-20, and 5 μl water. The reaction was incubated at 37 °C for 30 minutes in a thermomixer (ThermoMixer C Model 5382, Eppendorf) at 1000 RPM and cleaned up with Zymo DNA Clean and Concentrator-5 Kit. Purified DNA was eluted in 21 µl of elution buffer.

### PCR amplification, library quality control and concentration

All products of transposed fragments were pre-amplified using primers that included Illumina adaptors. The PCR reaction mixture was 2.5 μl 25 μM primer Ad1, 2.5 μl 25 μM primer Ad2 or i5/i7 primers (see Key resources table for oligo sequences), 25 μl 2x NEBNext Master Mix, 20 μl transposed sample; and the cycling conditions were as follows: 72 °C for 5 min, 98 °C for 30 sec, then 5 cycles of: 98°C for 10 sec, 63°C for 30 sec, 72°C for 1 min, and a hold at 4°C. To determine additional cycles, a qPCR using 5 μl of the pre-amplified mixture was performed as follows: Reaction mixture was 0.5 μl 25 μM Primer Ad1, 0.5 μl 25 μM Primer Ad2.x, 0.24 μl 25x SYBR Green (in DMSO), 5 μl 2x NEBNext Master Mix, 5 μl pre-amplified sample, 3.76 μl water; cycling conditions: 98 °C for 30 sec, then 20 cycles of: 98 °C for 10 sec, 63 °C for 30 sec, 72 °C for 1 min, and a hold at 4°C. The additional cycles performed to amplify the library were based on the one-third of the maximum fluorescent intensity calculated based on linear Rn versus cycle plot, as previously determined (Buenrostro et al., 2015a; Buenrostro et al., 2015b). A final amplification of the remaining 45 μl of pre-amplified DNA was run on the thermocycler with additional cycles determined as described above without additional reagents. Purification of the amplified library was performed using a Zymo DNA Clean and Concentrator-5 Kit and eluted in 20 μl water. All procedures to prepare the library were performed as described by (Corces et al., 2017). Library quality and fragment size distribution were assessed by High Sensitivity DNA ScreenTape assay D1000 kit (Agilent Technologies, Santa Clara, CA, USA), and the concentration was determined using a Qubit fluorometer (LifeTechnologies, Paisley, UK).

### Sequencing

To identify regions with open chromatin, we performed high-throughput DNA sequencing of the tagmented library on a NextSeq 500 System (Illumina) to generate paired-end, 75-cycle reads.

### ATAC-seq data analysis

We used the Cutadapt program (Martin, 2011) with options -a CTGTCTCTTATACACATCT -A AGATGTGTATAAGAGACAG -m 20 and -q 20 to remove the respective adapter sequences. We aligned the sequence reads to the mouse genome (assembly GRCm38 [mm10], Ensembl version 100) using BWA-MEM with the -M option (Li, 2013). We removed duplicates using the PICARD MarkDuplicates tool with the - REMOVE_DUPLICATES true option. We identified peaks using MACS2 callpeak with the --format BAMPE -g 1.87e9 –qvalue 0.05 option (Zhang et al., 2008). We used the soGGi package to identify consensus peaks and to remove peaks that overlapped with blacklisted regions (ENCFF547MET). We used the Rsubread package to count reads that occur within consensus peaks that exist in 2 or more biological replicates. We used the DESeq2 package (Love et al., 2014) to normalize counts and identify significant differentially accessible regions (adjusted P-value < 0.05). We used the ChIPseeker package (Yu et al., 2015) to annotate significant peaks with genomic features (e.g., promoter, 1^st^ intron, etc.). We used the GOstats package (Falcon and Gentleman, 2007) to test for statistical enrichment of GO terms for genes that had a promoter peak (within 1000 bp upstream and 500 bp downstream from a transcription start site) that was significantly greater in inf-μMT than conv-WT. Custom plots were generated within the R software environment (Team, 2022) and the ggplot2 package (Wickham, 2009). The ATAC-seq data is available from the ArrayExpress public repository (E-MTAB-10086).

### Chromatin immunoprecipitation (ChIP)

Bone marrow-derived macrophages from mice bone marrow were isolated by FACS as described above, and a ChIP assay was performed as previously described (Acevedo et al., 2007). ChIP was performed using mouse anti-STAT3 IgG1 antibody (Santa Cruz). DNA was quantified using qPCR with the following primer pairs: IL-10 promoter binding site for Stat3 Forward: 5′-TCATGCTGGGATCTGAGCTTCT-3′, Reverse: 5′CGGAAGTCACCTTAGCACTCAGT-3′ as previously described (Lucas et al., 2005). The following primers for r18S were used as a non-specific primer control: Forward: 5’- CATTCGAACGTCTGCCCTATC-3’, Reverse: 5’-CCTGCTGCCTTCCTTGGA-3’. qPCR values were normalized as percent recovery of the input DNA and expressed as % of input.

### Image analysis

ImageJ 1.53c software (National Institutes of Health, USA) and the Fiji image processing package (Schindelin et al., 2012) were used to analyze confocal fluorescent images. Leica Application Suite X version 3.5.6.21594 was used to process images acquired in a tile scan mode.

### Statistical analysis

All data were analyzed using GraphPad Prism version 8.0 (GraphPad Software, San Diego, CA). Statistical significance was determined by one-way or two-way ANOVA followed by Tukey’s multiple comparisons test or two-tailed, unpaired Student’s t-test. A statistically significant difference was defined by P-values<0.05. The data represent means ± SEM. The experiments were repeated at least twice.

### ATAC-seq data

The ATAC-seq data is available from the ArrayExpress public repository (E-MTAB-10086).

## Author contributions

Conceptualization: D.T.G., E.K.J., R.T.G.

Data curation: L.S.S., D.C.

Formal Analysis: L.S.S., D.C.

Funding acquisition: D.T.G. and K.A.F

Investigation: L.S.S., C.S.F., Y.N., J.N.C., G.H., N.P.T.

Methodology: L.S.S., D.T.G., B.M., E.K.J., R.T.G., E.L., Z.A., E.S.

Project administration: D.T.G.

Resources: D.T.G., Z.A., E.L.

Software: D.C., T.R.

Supervision: D.T.G.

Validation: D.T.G., E.K.J., K.A.F., R.T.G.

Visualization: L.S.S.

Writing original draft: L.S.S.

Writing review & editing: D.T.G., L.S.S., E.K.J., R.T.G., K.A.F.

## Acknowledgments

We thank the Microscopy Core Facility of the Medical Faculty at the University of Bonn for providing support and instrumentation. The authors also acknowledge Dr. Joanne Thompson from the University of Edinburgh and the European Malaria Reagent Repository (www.malariaresearch.eu) as the source of the GFP-expressing *P. chabaudi*. In addition, we thank the staff of the Electron Microscopy Core Facility at the UMass Chan Medical School.

## Supporting information

**Figure S1 – Experimental design**. WT and uMT^−/−^ mice were injected intraperitoneally with 10^5^ erythrocytes infected with *Plasmodium chabaudi chabaudi* AS-GFP. The control groups were uninfected WT and uMT^−/−^ mice injected intraperitoneally with an equivalent volume of vehicle (1xPBS). Parasitemia was assessed every seven days by flow cytometry (see S2A Fig). Analysis was based on *Plasmodium chabaudi* phases: acute (black bar) first peak of parasitemia, chronic (blue bar) sub-patent parasitemia, and convalescence (yellow bar) infected for the WT group that cleared the parasite (PCR negative). Most of the analyses were performed when infected WT mice were convalescent.

**Figure S2. Flow cytometry and PCR analyses of parasitemia in mice.** (A) Gating strategy to assess parasitemia in mice based on TER119 expressed by red blood cells and GFP expressed by *Plasmodium chabaudi*. Blood samples were fixed in 0.025% glutaraldehyde in 1XPBS and stained with anti-TER119-PE-Cy7. The *Plasmodium chabaudi chabaudi* AS strain expressing GFP was used to identify infected RBCs. Four sequential gates were applied to exclude debris (upper left) and doublets (upper right), to include RBCs (bottom right) and to include GFP+ events (bottom left). Infected RBCs are identified as TER119+GFP+ events. (B) Nested PCR based on *Plasmodium* spp. chromosome 12 has a high sensitivity to detect parasite DNA in whole blood samples from mice. The gel shows nested PCR products performed on serial dilutions (from 1:10 to 1:10^6^) of *P*. *chabaudi*-infected blood. Lane “L” contains the 100-bp DNA ladder; the second and third lanes are products from nested PCR of *P*. *chabaudi*-infected blood samples with 0.5% and 0.1% parasitemia, respectively; fourth through ninth lanes are a logarithmic serial dilution from infected blood (initial parasitemia of 0.1%). Blood from uninfected mice was used as the diluent and as a negative control (last lane on the right). Image is representative of two independent experiments.

**Figure S3. Parasitemia levels of mice used for metabolic cage analysis.** (A) Parasitemia (days 8, 30, and 50 post-infection) from C57BL/6J WT (closed black circle) and μMT^−/−^ (red triangle) mice infected with fluorescent *P. chabaudi* AS expressing GFP are shown as a percentage of TER119+GFP+ events. Blood samples of uninfected C57BL/6J WT (white square) and μMT^−/−^ (blue diamond) mice were used as negative controls (n=3 for all groups). (B) PCR amplification of blood samples from uninfected WT (uninf-WT), uninfected μMT^−/−^ (uninf-μMT), infected convalescent WT (conv-WT), and infected μMT^−/−^ (inf-μMT) mice at 50 dpi, n=3.

**Figure S4. Chronically infected μMT^−/−^ mice had reduced dendritic cells (DCs) and increased macrophages and hemozoin accumulation in the spleen.** Confocal microscopy images from spleen sections antibody stained with anti-F4/80 (green), anti-CD11c (magenta), and hemozoin (yellow) and examined by confocal reflection from uninfected WT (uninf-WT), uninfected μMT^−/−^ (uninf-μMT), convalescent WT (conv-WT), and infected μMT^−/−^ (inf-μMT) mice 99 dpi (left panel). XZ and YZ confocal plane from infected μMT^−/−^ mice showing colocalization between F4/80, CD11c, and hemozoin (macrophage); and CD11c with hemozoin (dendritic cells) at 99 dpi (right panel), bar = 50 μm. Bar graphs showing the quantification of A. Data are shown as mean ± SEM. * P<0.05, ** P<0.01, n.s. = not significant, as determined by one-way ANOVA followed by Tukey’s multiple comparisons test.

**Figure S5. High circulating levels of IL-10 in infected μMT^−/−^ mice are not produced by CD4-, CD4+, or CD14+ cells from the peripheral blood.** Blood samples from uninfected WT (uninf-WT), uninfected μMT^−/−^ (uninf-μMT), convalescent infected WT (conv-WT), and infected μMT^−/−^ (inf-μMT) mice 97 dpi were treated with 20 ng/mL PMA, 1 μg/mL ionomycin, and 5 μg/mL Brefeldin A for 6 h. Red blood cells were lysed and IL-10 production from CD4-(A), CD4+ (B), or CD14+ (C) cells were analyzed by flow cytometry.

**Figure S6. IL-10 protein levels in total spleen lysate.** Spleen tissue samples from uninfected WT (uninf-WT), uninfected μMT^−/−^ (uninf-μMT), convalescent WT (conv-WT), and infected μMT^−/−^ (inf-μMT) mice were collected at 90 dpi, and levels of IL-10 were accessed by ELISA and expressed as pg/g of tissue, (n=5). Data are shown as mean ± SEM. ** P<0.01, *** P<0.001, as determined by one-way ANOVA followed by Tukey’s multiple comparisons test.

**Figure S7. Schematic diagram of IL-10R blockage in chronically infected μMT^−/−^ mice.** IL-10R blockade was achieved by injections of four doses of anti-IL-10R antibody and compared to injections with an isotype control. For the first two doses, 500 µg of antibody was administered per mouse; for the last two doses, 250 µg of antibody were administered per mouse. The same doses of an IgG isotype control were injected in the control group. Blood samples for CBC, CRP, or cytokine assays were collected before antibody injection (day 0) and on days 7, 13, and 21 post-antibody injections. Parasitemia and body weight were evaluated at days 0, 4, 7, 11,13, and 17 post antibody injection, and spleen tissue was collected at day 21 for histology.

**Table S1. Genomic coordinates from genes with significant differences in chromatin accessibility.**

**Table S2. Key resources table.** Table of critical reagents, organisms and strains, software and algorithms, instrumentation, and source of deposited data presented in this study.

